# Circuit reorganization in the *Drosophila* mushroom body calyx accompanies memory consolidation

**DOI:** 10.1101/2020.08.20.259135

**Authors:** Lothar Baltruschat, Philipp Ranft, Luigi Prisco, J. Scott Lauritzen, André Fiala, Davi D. Bock, Gaia Tavosanis

## Abstract

The capacity of utilizing past experience to guide future action is a fundamental and conserved function of the nervous system. Associative memory formation initiated by the coincident detection of a conditioned stimulus (CS, e.g. odour) and an unconditioned stimulus (US, e.g. sugar reward) can lead to a short-lived memory trace (STM) within distinct circuits [1-5]. Memories can be consolidated into long-term memories (LTM) through processes that are not fully understood, but depend on *de-novo* protein synthesis [6, 7], require structural modifications within the involved neuronal circuits and might lead to the recruitment of additional ones [8-17]. Compared to modulation of existing connections, the reorganization of circuits affords the unique possibility of sampling for potential new partners [18-20]. Nonetheless, only few examples of rewiring associated with learning have been established thus far [14, 21-24]. Here, we report that memory consolidation is associated with the structural and functional reorganization of an identified circuit in the adult fly brain. The formation and retrieval of olfactory associative memories in *Drosophila* requires the mushroom body (MB) [25]. We identified the individual synapses of olfactory projection neurons (PNs) that deliver a conditioned odour to the MB and reconstructed the complexity of the microcircuit they form. Combining behavioural experiments with high-resolution microscopy and functional imaging, we demonstrated that the consolidation of appetitive olfactory memories closely correlates with an increase in the number of synaptic complexes formed by the PNs that deliver the conditioned stimulus and their postsynaptic partners. These structural changes result in additional functional synaptic connections.

## Results

To start addressing systematically the mechanisms that support memory consolidation, we sought to investigate the properties of identifiable synapses in the MB of the adult brain of *Drosophila* after the establishment of LTM. Within the main MB input compartment, the calyx (MBC), PNs deliver olfactory information through cholinergic synapses to the intrinsic MB neurons, the Kenyon cells (KCs) (Figure 1A). In the MBC, large olfactory PN boutons are enwrapped by the claw-like dendrite termini of ∼11 KCs on average [26, 27], thereby forming characteristic synaptic complexes, the microglomeruli (MGs) [28] (Figure 1A-E), which display functional and structural plasticity in adaptation and upon silencing [29-31]. The pheromone and odorant 11-cis-Vaccenylacetate (cVA) activates specifically those PNs that project their dendrites to the DA1 glomerulus in the antennal lobe (AL) [32-34] (Figure 1A, B; DA1-PNs). The DA1 glomerulus is mostly excluded from complex processing of sensory information in the AL [35, 36], suggesting that by genetically marking the DA1 PNs we could identify the individual boutons in the MBC that deliver the olfactory response to cVA. We tested this by recording with volumetric calcium imaging the response to odour stimulation in the MBC of animals expressing a genetically encoded calcium indicator tethered at the KC postsynapses (*MB247-homer::GCaMP3*) [30] in combination with a presynaptic fluorescent tag (*UAS-td-Tomato*) expressed in DA1-PNs only (Figure 1C). Regions of interest (ROIs) containing fluorescently labelled DA1-MGs showed a postsynaptic response specifically tuned to cVA stimulation (84% of the fluorescently labelled MGs responded to cVA only; n=7 animals, 72 boutons; Figure 1C). Taken together, by selecting the combination of the cVA odorant and the DA1 subset of PNs we established fly lines in which we can track a fly’s neuronal response towards a specific odour on the level of individual synaptic complexes in the MBC (Figure 1A-C).

**Figure 1.**
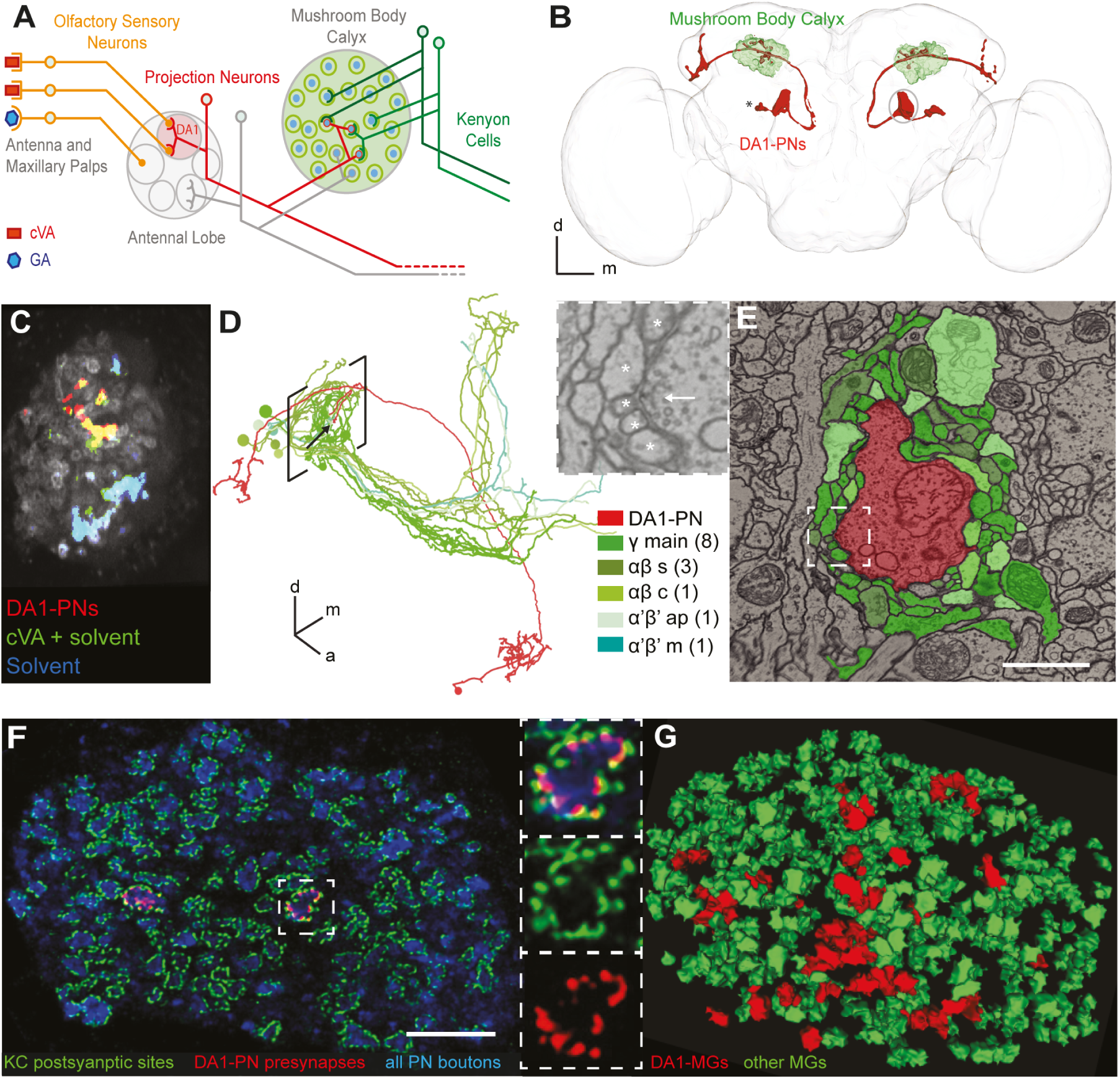
Identification of the synapses in the MBC responding to cVA odour stimulation. **(A)** Schematic representation of the olfactory circuit starting from the activation of specific Olfactory Sensory Neurons (OSNs) by two exemplary odours cVA and GA. In the AL cVA-responsive OSNs converge to the DA1 glomerulus (pale red), where they synapse onto DA1-PNs (red). These deliver the cVA signal to the MBC via axon collaterals that terminate with boutons forming large synaptic complexes, the MGs. Postsynaptic KCs are represented in green. **(B)** Reconstruction from a full confocal serial section set of the DA1-PNs (red; *R37H08-Gal4> UAS GAP43::Venus*); MBC (green; *MB247-Dα7::GFP*); DA1-PN cell bodies (*); brain neuropil (light grey; α-synapsin antibody). **(C)** Volumetric calcium imaging of the calyx of flies carrying *MB247-Homer::GCaMP3* (grey) and in which DA1-PNs are genetically labelled (red; *R37H08-Gal4> UAS tdTomato*). cVA-elicited postsynaptic responses (green) are specific to DA1-PNs, confirmed by the overlap between the two channels (yellow). Generic response to the solvent (cyan= overlap of the responses to cVA+ EtOH, green, and to EtOH only, blue). **(D)** Single DA1-PN (red) and the 14 KCs (green) postsynaptic to the DA1-PN bouton indicated by the arrow. Tracings performed on the EM FAFB dataset [37]. Square brackets indicate location of MBC. Different green shades represent different KC subtypes (as in E). Numbers in brackets in the legend represent the number of cells. **(E)** Single EM section through the MG (arrow in D); scale bar = 1 µm. White square is magnified in left top panel with arrow pointing to a T-bar of the AZ and * labelling fine dendritic postsynaptic profiles of KCs. **(F)** Single plane confocal image of the MBC displaying PN boutons (blue; α-synapsin antibodies); the PSDs of KCs (green; *MB247-Dα7::GFP*) and the AZs of DA1-PN boutons only (red; *R37H08-Gal4 > UAS-brp-short*^*cherry*^) identifying the cVA-responsive MGs. Scale bar = 10 µm. The MG in the white square is magnified in the right panels. **(G)** Automated 3D reconstruction of a confocal stack, including the image shown in (F). The reconstruction of MGs is based on Dα7-GFP (green) (see also Figure S1) and MGs receiving presynaptic input from DA1-PNs are marked by Brp–short^cherry^ (red). All other MGs are in green. Full genotypes used and statistics for all Figures are included in the Supplemental Genotypes and Statistics table.

To gain insight into the complexity of the MG microcircuit formed by a single DA1-PN bouton we took advantage of the availability of a whole brain electron microscopy (EM) volume of an adult female fly (FAFB) [37]. In this set of data we reconstructed a complete MG connectome by tracing neurites from every pre- and postsynaptic contact of a DA1-PN bouton until the corresponding neuron’s identity was anatomically determinable (Figure 1D, E; Supplementary Table 1). This particular DA1-PN bouton made 33 excitatory, cholinergic contacts, all polyadic and identifiable by the presence of a T-bar and of a synaptic cleft (Figure 1E, inset), apposed to 277 postsynaptic profiles. Most profiles (248) postsynaptic to the bouton originated from 14 KCs of 5 different subtypes: γmain (8), αβs (3), αβc (1), α’β’ap (1) and α’β’m (1) [38](Figure 1D, E; Supplementary Table 1, Movie 1). γmain profiles were thus the most abundant in this particular bouton, although DA1-PN boutons are located within a region of the MBC predominantly occupied by αβs KCs [39]. Each KC contacted the bouton with a single claw receiving 8 to 25 presynaptic inputs from the PN bouton, in line with previous estimates [26, 31]. Within the MG, the bouton received presynaptic input from 4 cells: two additional γmain KCs forming divergent triads that included a KC, the PN bouton and the anterior paired lateral neuron (APL) [40]; APL itself and one of the two Mushroom Body Calyx 1 neurons (MB-C1) (Supplementary Table 1 and described elsewhere). Taken together, a single MG represents a highly complex microcircuit, involving many neurons (19 in this example) of different types (here 8).

To investigate whether such a complex structure could undergo plastic changes, we designed a setup to reveal and measure the properties of identifiable MGs following olfactory conditioning.

In confocal images, we highlighted cVA responsive MGs in the MBC by expressing only in DA1-PNs the presynaptic active zone (AZ) marker Brp-short^cherry^ (Figure 1F) [29, 41]. The postsynaptic densities (PSDs) of KC dendrites were decorated by cell-type specific genetic expression of a GFP-tagged Dα7 subunit of the acetylcholine receptor (Figure 1F) [29]. MGs could thus be identified with a software-based automated 3D-reconstruction tool exploiting the *MB247-Dα7::GFP* signal and were then classified as DA1-PN positive if they additionally displayed Brp-short^cherry^ co-labelling (DA1-MG; Figure 1G; Figure S1). Further, we established an appetitive associative conditioning paradigm using cVA or geranyl acetate (GA) as CS in STM or LTM paradigms (Figure 2A; Figure S2A-C; see Methods) and applied it to flies expressing the reporters mentioned above (Figure 2B, F, J). Alternatively, we mock-trained the flies by presenting each odour or the sugar reward at 2 min intervals to avoid the formation of appetitive association (Figure 2A, B, F, J) [42]. GA was chosen as it activates a separate and non-overlapping set of PNs in comparison to cVA [43] and 5% EtOH was added to provide a food-related context to the starved flies (Figure S2B) [36, 44]. To assess if MGs formed by DA1-PN boutons (DA1-MGs) underwent morphological modifications after learning, we prepared for confocal imaging female fly brains dissected at 1 min (STM) or at 24 h (LTM) after training. After STM establishment (Figure 2B) the total number, MG volume and lumen volume of DA1-MGs was unchanged in cVA conditioned (cVA CS+) flies compared to the GA conditioned (GA CS+) or mock control groups (average MG numbers: mock 28.91; GA CS+ 30.40; cVA CS+ 28.76; n= 10-17; Figure 2C-E). However, in the LTM paradigm (Figure 2F) the MG volume and the lumen volume of DA1-PN MGs were decreased in cVA CS+ flies compared to GA CS+ or to mock-control flies (Figure 2G, H). In addition, the total number of DA1-MGs was increased (average MG numbers: mock 27.31; GA CS+ 27.47; cVA CS+ 32.06; n= 18-32; Figure 2I). Thus, LTM, but not STM, was accompanied by an input-specific structural reorganization of the MBC circuit, including an increase in MGs number. These changes were specific to the conditioned odour, as they did not appear in the DA1-MGs when the conditioned odour was GA. These data suggest that the neurons delivering the CS form new boutons, which are of smaller size and enveloped by KC claws.

**Figure 2.**
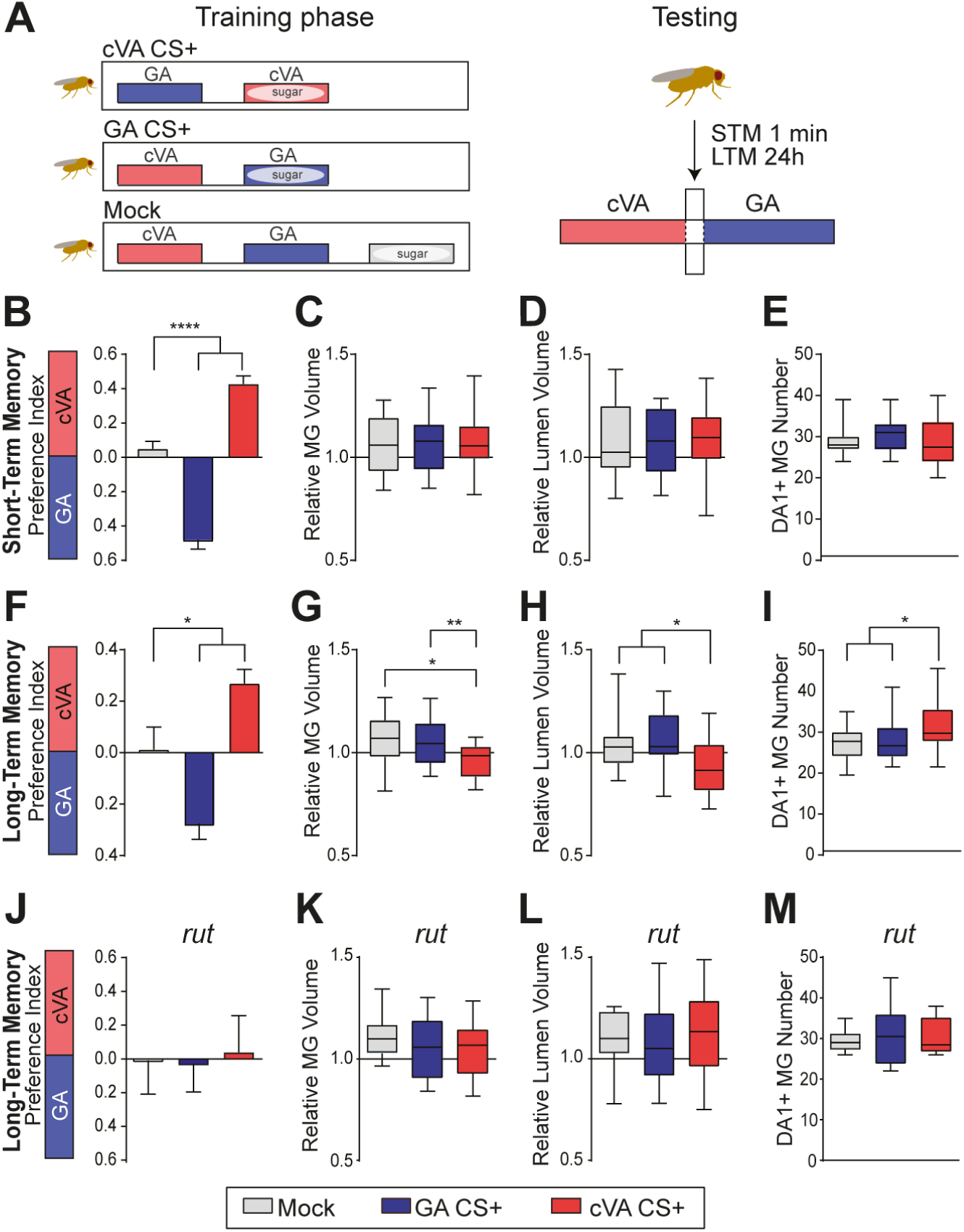
Microglomeruli undergo structural changes upon appetitive long-term memory formation. **(A)** Schematic illustration of the appetitive conditioning paradigm. For training, the conditioned odour cVA (red box) or GA (blue box) is paired with sugar. In STM experiments flies are trained for 2 min with a 2 min interval between CS+ and CS-presentation and tested 1 min after training. In LTM experiments flies are trained for 5 min + 5 min with a 2 min stimulus interval and are tested 24 h after training. In the mock control the two odours and the sugar reward are presented in a temporally spaced sequence with a 2 min inter-stimulus pause. **(B, F, J)** Preference indices of flies *R37H08-Gal4/MB247-Dα7::GFP, UAS-brp-short*^*cherry*^ in the STM (n = 19 – 25) **(B)** or in the LTM (n = 14 - 19) paradigm **(F)** and preference index of *rut* mutant flies in LTM **(J)**, (*p* > 0.05, n = 17-18). Preference index values of the mock control group (grey bars) were compared to groups trained with GA CS+ (blue bar) or cVA CS+ (red bar). Multiple comparisons are tested throughout this study with one-way ANOVA with Bonferroni correction. Significance level is set at p < 0.05. *p < 0.05, ****p < 0.0001. In this and all other bar graphs, bars and error bars indicate mean and SEM, respectively. **(C, G, K)** MG volume is defined by the area enwrapped by the postsynaptic densities of KC dendrites. MG lumen volume **(D, H, L)** is the volume of the lumen as defined in Figure 1F. In STM the relative volume of DA1-MGs **(C)** and of their lumen **(D)** is not different between groups (p > 0.05; n = 15-20). In LTM the MG volume **(G)** and the lumen volume **(H)** of DA1-MGs in flies trained with cVA CS+ are smaller than in flies from the mock control group or of flies trained with GA CS+ (* p < 0.05, **p < 0.01; *n* = 19–25). **(E, I, M)** Number of DA1-PN positive MGs is unaffected in STM **(E)** (*p* > 0.05; n = 18 – 24). In LTM, number of DA1-PN positive MGs in cVA CS+ trained flies is higher compared to flies of the mock control or GA CS+ group **(I)** (*p < 0.05; n = 18-24). **(K-L)** The structural modifications of DA1-MGs in cVA CS+ trained flies after the appetitive LTM protocol were suppressed in *rut* mutants (*p* > 0.05, n = 13–21). In all box plots in this and the following figures, the edges of the boxes are the first and third quartiles, thick lines mark the medians, and whiskers represent data range.

Olfactory associative learning relies on the function of the Ca^2+^/CaM-dependent adenylyl cyclase Rutabaga [42, 45, 46] and a defining trait of LTM is its dependence on protein synthesis [6, 47, 48]. Indeed, a mutation in the *rutabaga* gene (*rut*^*2080*^) [49] or feeding flies with the protein synthesis inhibitor cycloheximide (CHX) immediately after training abolished LTM (Figure 2J; Figure S2D). Importantly, loss of *rut* function or CHX feeding also suppressed the structural changes in the DA1-MGs supporting the correlation between LTM formation and structural changes in the circuit (Figure 2K-M; Figure S2E-G).

The increase in DA1-PN positive MG number after LTM formation with cVA CS+ suggests that new boutons might be formed during consolidation. To gain insight into the cellular fundamentals of these modifications, we expressed in DA1-PN axons the membrane-tagged fluorescent protein *UAS-GAP43::Venus* together with *UAS-brp-short*^*cherry*^ and highlighted the postsynaptic densities on KC dendrites using *MB247-Dα7::GFP* (Figure 3A-D). Serial optical sections of the MBCs of these flies trained with cVA CS+, with GA CS+ or in the mock paradigm (Figure S2H) were used to generate 3D reconstructions that were then aligned to a reference brain (JFRC2 [50]). The DA1-PN axons were then traced in the aligned high-resolution scans of the MBC (Figure 3E). The DA1-PN boutons were highly clustered in the dorsal-posterior part of the calyx [51], supporting the view that the localization within the MBC of DA1-PN boutons is not entirely random (Figure 3F) [52]. The total length of DA1-PN collaterals measured from the point where they leave the inner antennocerebral tract (iACT) was increased in flies that had formed LTM after cVA CS+ training compared to mock-control flies (Figure 3G). In addition, the total fraction of the MBC volume containing DA1-PN positive boutons was increased in flies that had formed cVA CS+ LTM (Figure 3H). These observations suggest that during consolidation, additional boutons are created by local growth at existing DA1-PN collaterals (Figure 3I).

**Figure 3.**
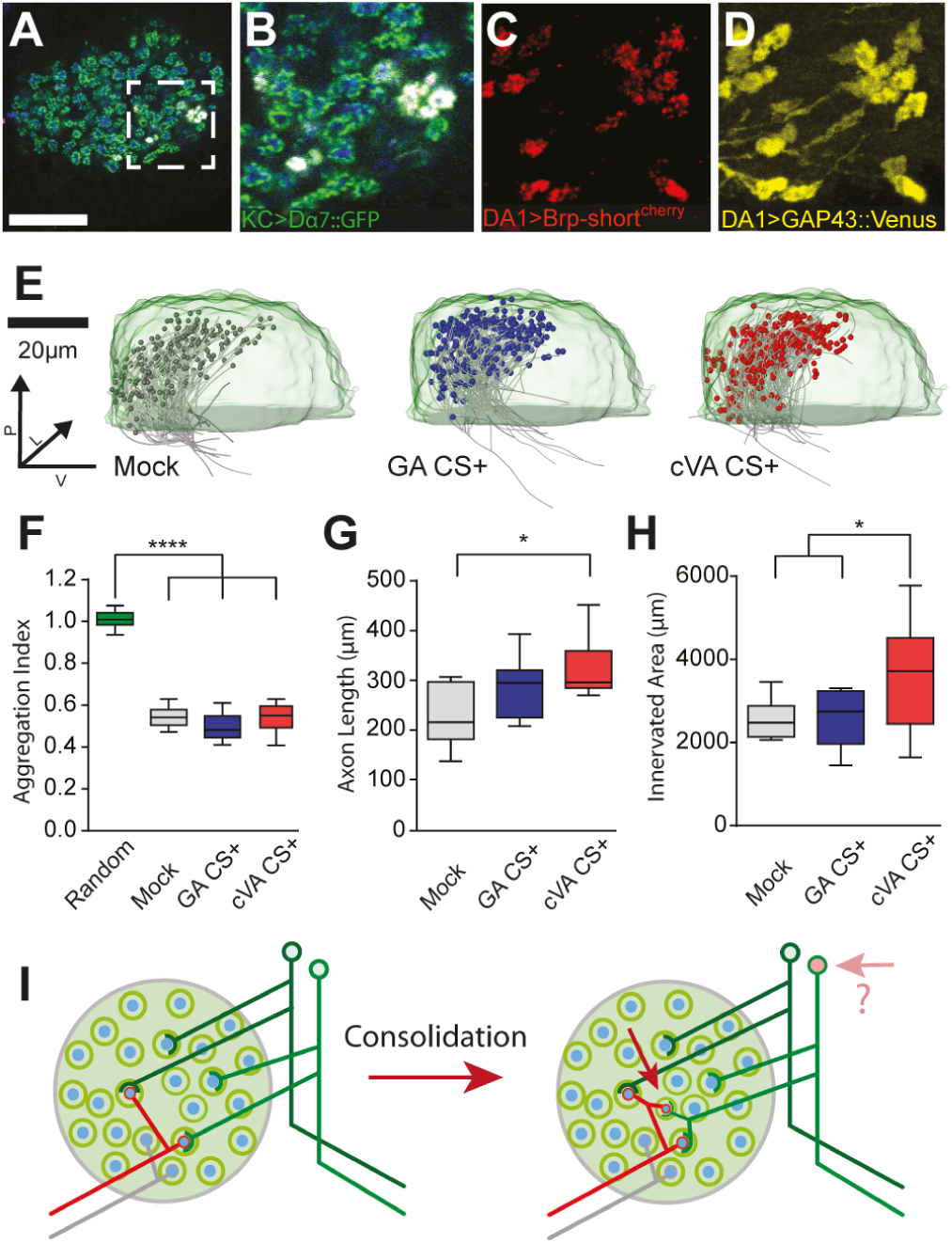
Modifications of axon collaterals and wiring properties of projection neurons within the mushroom body calyx after long-term memory formation. **(A)** Single optical section of the MBC of flies expressing Dα7GFP (green) in the KCs and Brp-short^cherry^ (red) plus GAP43-Venus (yellow) in DA1-PNs (*R37H08-Gal4*); PN boutons (blue; anti-Synapsin antibodies). **(B-D)** Magnification of the white square in (A) displaying the merge as in A **(B)** or a maximum-intensity projection of Brp-short^cherry^ **(C)** or of GAP43-Venus **(D)** signals. **(E)** Medial view of registered PN axons (grey) with traced boutons (grey, blue, red spheres) within a standard calyx (light green). The registered PN traces are of mock (grey), GA CS+ (blue) or cVA CS+ (red) trained groups. n = 10 for each group. **(F)** Boutons are highly clustered independently of the treatment (Clark and Evans Aggregation index compared to a hypothetical random distribution; ****p < 0.0001). **(G)** Total collateral axons length of mock control, GA CS+ or cVA CS+ flies. (*p < 0.05; n = 10). **(H)** The convex hull volume containing all DA1-boutons in the MBC per condition is increased in cVA CS+ flies compared to mock control and GA CS+ group. **(I)** We suggest that the increased number of MGs after consolidation is due to the formation of additional boutons responding to cVA. The additional boutons form full MGs, as postsynaptic profiles of KCs surround them. It is unclear whether this reorganization might lead to the recruitment of additional responding KCs (see Discussion).

To address whether the observed structural changes within the MGs after LTM impact on the functional representation of the CS in the MBC we analysed calcium dynamics in KC dendrites. For this, we utilized flies carrying *MB247-homer::GCaMP3* [30] in combination with volumetric calcium imaging (Figure 4A). We utilized this simple genotype to guarantee that flies performed well in LTM experiments (Figure S2I). We measured calcium response in the entire MBC volume during a single odour application (5 s odour stimulation) of either cVA (1:400 in 5% EtOH) or EtOH alone (5%). To identify areas with increased calcium dynamics during odour stimulation we overlaid a grid consisting of 5×5 µm^2^ ROIs over each optical section of the volumetric time series. Based on the grid segmentation, we then calculated the average ΔF/F% for each ROI in the MBC. ROIs were classified as odour responsive if the measured calcium response exceeded a set threshold (ΔF/F% > 3x standard deviation) during the first 2 s of stimulation (Figure 4B, C; Figure S3A, B). The response pattern elicited specifically by cVA was defined after subtraction of the EtOH response (Figure 4C; Figure S3C, D; see also Figure 1C). This approach yielded a comprehensive picture of the functional odour response over time in the MBC of trained or control flies (Figure S3D, E). After appetitive LTM formation, the percentage of cVA-responsive ROIs was increased in cVA CS+ flies compared to the mock control (Figure 4D) (n = 7, p < 0.05), suggesting that the additional DA1-PN boutons are functionally connected to their postsynaptic KC counterparts and capable of initiating a response in the postsynaptic KCs. We further analysed the calcium dynamics over time of all animals of the mock or cVA CS+ group (Figure 4E, F). Linear regression analysis of the fluorescence change during odour stimulation showed a steeper drop of the linear fit in cVA CS+ flies (R^2^ = 0.6429) towards baseline compared to flies of the mock control group (R^2^ = 0.1124). Besides, the response towards the odour was more variable in mock-trained flies compared to the cVA CS+ flies (Figure S3F). Initially (0-4s after start of stimulation), the total response towards cVA stimulation was indistinguishable between mock control and cVA CS+ group. However, at subsequent time points (4-7s after start of stimulation) responses were significantly lower in KC dendrites of the cVA CS+ group compared to the mock control, showing a faster calcium decay towards the trained odour in CS+ flies (n = 7, p < 0.05) (Figure S3G). Together, these data indicate a temporal sharpening of the odour response.

**Figure 4.**
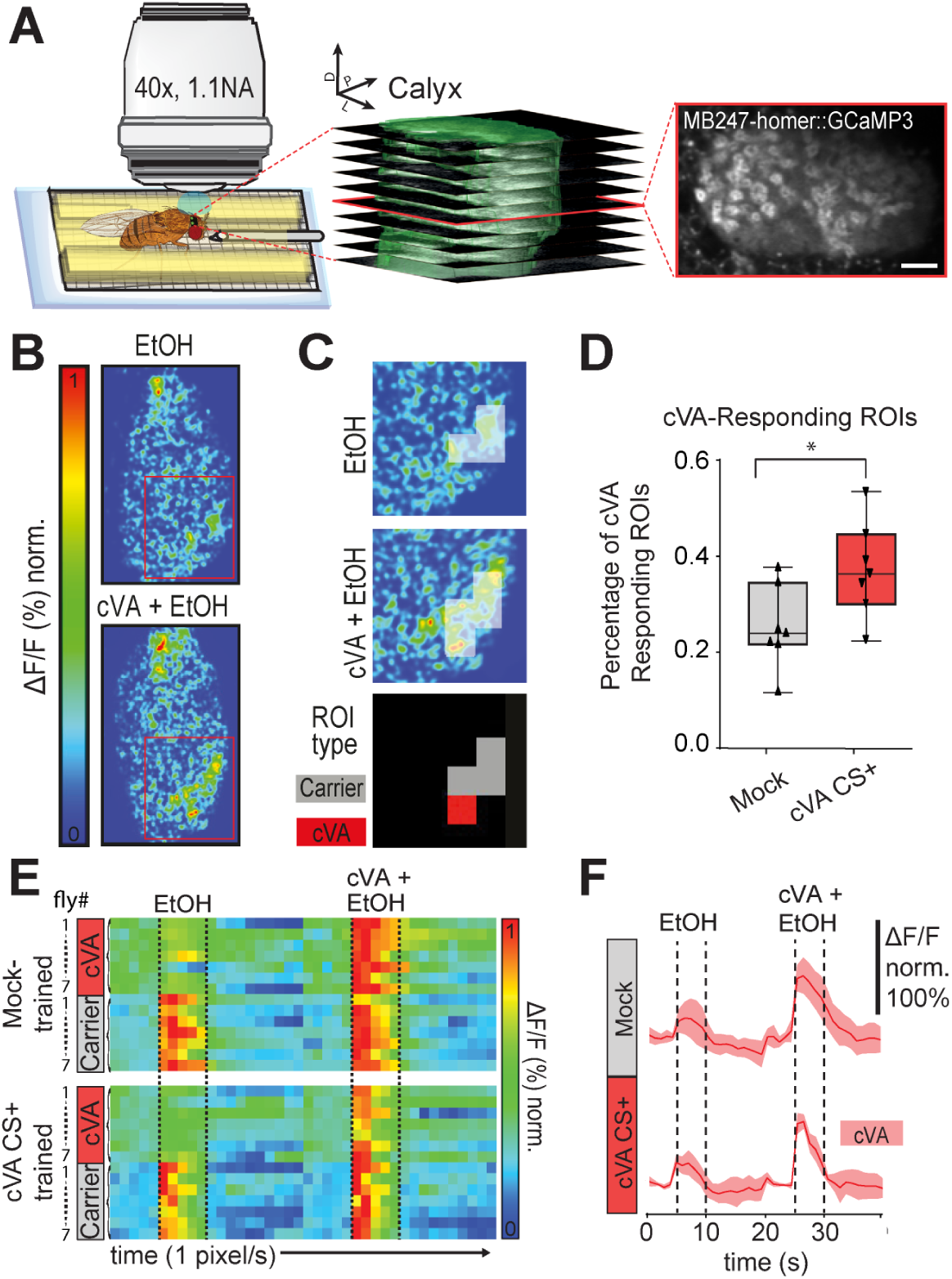
Long-term memory-induced functional plasticity in the mushroom body calyx. **(A)** Two-photon *in vivo* imaging setup. Schematic of a fly placed on a custom-made holder under a two-photon microscope equipped with a 40x 1.1 NA water immersion objective. The odour is delivered for 5s with a moisturized, constant air stream through a 1.2mm cannula. (Central panel) Z series of the entire MBC volume of flies expressing post-synapse-tagged Homer-GCaMP3 imaged during odour application at 1Hz (10 optical sections per volume, 4µm step size). (Right panel) A single slice of the image stack shown in the middle panel. Scale bar = 10 µm. **(B)** Representative optical section from a volumetric time series showing false-coloured response of KC dendrites to 5 s exposure to EtOH (top) or cVA + EtOH (bottom). **(C)** Magnification of the red square area in (B). 5×5 µm^2^ ROIs were classified as cVA-responsive (red) if they were only active during cVA + EtOH application, but did not respond to EtOH alone. **(D)** The percentage of cVA responsive ROIs increased after LTM acquisition compared to the mock control (*p < 0.05; n = 7). **(E)** Dynamics of ΔF/F% changes over time in KCs of *MB247-homer::GCaMP3* flies after mock training (top) or LTM acquisition (bottom). Each row of the heat map represents average responses per animal of all cVA responsive ROIs (pink) or of all carrier EtOH responsive ROIs (grey) within one MBC over time. Each column represents one 1s. Flies are first exposed to the EtOH (5s) and then to cVA + EtOH (5s) as indicated by the dashed lines. **(F)** Plot of average calcium dynamics over time of cVA responsive ROIs during 5 s stimulation with EtOH or with cVA in EtOH (dashed lines) in mock-trained (top) or cVA CS+ (bottom) flies (n = 7 per condition).

## Discussion

We report input-specific reorganization of the adult MBC circuit associated with the formation of long-term appetitive memory. By visualizing presynaptic markers in PNs and the KC postsynaptic densities, we uncover an increase in the number of PN boutons and at the same time reveal that these boutons are enveloped by KC postsynaptic profiles, suggesting that new MGs are formed during memory consolidation. These findings are particularly remarkable, given the high degree of complexity of the MG microcircuits revealed by our EM reconstruction and including the dendrite claws of multiple KCs of distinct subtypes. The cellular mechanisms leading to the increased number of odour-specific complex MGs remain to be clarified, but they will require a tight coordination between pre- and postsynaptic partners. We suggest that they could be driven by intrinsic reactivation of KCs during the consolidation phase [53, 54] or by modulatory inputs into the calyx [38, 55-58]. In either case, we expect a complex pattern of activation that might be difficult to reproduce in artificial settings [29, 59]. In agreement with our present report, the density of PN boutons in the MBC increases after appetitive long-term conditioning in honeybees and transiently also after avoidance learning in leaf-cutting ants [60, 61]. In comparison to those data, using genetic and functional identification of PN subsets we reveal here that the structural modifications are specific and limited to the PNs conveying the conditioned odour. Importantly, our *in vivo* functional imaging data support the view that the circuit reorganization that we describe leads to additional functional MGs responding to the conditioned odour. Additionally, they demonstrate a specific change in functional response in the KC dendrites towards the trained odour as the calcium levels drop faster towards baseline after appetitive associative conditioning. The faster decay kinetics could contribute to a more efficient temporal summation of responses or refine the KC response [62]. They are in line with recent data from appetitive conditioning experiments from the honeybee possibly rooted in inhibitory modifications [63]. An important open question is the effect of the increased number of responding MGs on the pattern of KC activation. KCs respond sparsely to odour input and require the coincident activation of multiple of their claws to produce an action potential [64]. Our data might underlie the addition of connections between the active PNs and, preferentially, a set of already responding KCs, leading to facilitated response to the conditioned odour, but without changing the set of responding KCs. A recent publication suggests an exciting alternative view. After aversive LTM establishment, the number of KCs responding to the conditioned odour is increased [65]. If we hypothesize that appetitive conditioning could lead to a similar outcome, our data could provide anatomical and functional support to these findings. The combination of these data suggests that the pattern of KC response is modulated by experience during the adult life of the animal and might represent a rich signifier of sensory stimulus and context.

## Acknowledgements

We thank K. Keppler, M. Schoelling, A. Mueller and J. C. Vijayakumar for technical assistance. We are grateful to C.B. Fisher, S.A. Calle-Schuler, N. Sharifi, B. Gorko, L. Kmecova, I.J. Ali, N. Masoodpanah, J. Hsu and F. Li in D. D. Bock’s lab for their support in tracing evaluation. We thank Y. Aso, HHMI Janelia, the Kyoto Drosophila Genetic Resource Center and the Bloomington Stock Center for fly lines and FlyBase. We are grateful for their assistance to C. Moehl in the establishment of the cluster analysis, S. Dipt with initial calcium imaging experiments and R. Court for providing the brain aligner and support. We thank S. Sachse, D. Isbrandt, T. Hige, S. Remy, M. Nawrot, O. Barnstedt, the members of the Tavosanis lab for discussion and/or critical reading of the manuscript and B. Schaffran for help with video editing. L.B. acknowledges support by the Bonn International Graduate School of Neuroscience. This work was supported by the DFG FOR 2705 to G.T.

## Author Contributions

L.B. and G.T. conceived the project and designed the experiments. L.B., P.R. and L.P. constructed fly strains, performed behavioural experiments, produced and analysed the anatomical data. J.S.L. and D.D.B. established the set of EM data and P.R. performed the tracings presented here. Scripts and routines for the analysis were established by L.B. and L.P. Functional imaging experiments were designed and performed by L.B and L.P. with support by A.F. The manuscript was written by G.T, L.B., P.R and L.P.

## Methods

### EM reconstruction and identification

Neuron skeletons were reconstructed in a serial section transmission electron microscope (ssTEM) volume of a complete female adult *Drosophila melanogaster* brain [37] and manually traced using CATMAID [66]. Thus, traced neuron skeletons represent the branching of neurons and the location of their cell bodies and synapses. Chemical synapses were manually annotated and identified consistently with the criteria of other CATMAID-based *Drosophila* connectomic studies [37]: 1) an active zone (AZ) surrounded by vesicles, 2) a presynaptic specialization (*e*.*g*. T-bar), 3) synaptic cleft and 4) a post synaptic density zone (PSD), which however can be absent. If the PSD is absent, we annotated all cells along the synaptic cleft as postsynaptic [37, 67]. Neuron identity is based on previously described morphologies in light microscopy (KC subtypes, APL, MB-C1, PN), such as dendritic branching, axonal projection and location in the neuropil [38, 40, 52, 68, 69]. Additionally, we performed a neuron search against a light microscopy dataset in NBLAST [70], as described in [37] for PN subtype identification. 3D reconstructions of the PN bouton and KC claws from ssTEM sections were created manually with the ImageJ plugin TrakEM2 [71].

### Fly strains

Flies were raised at 25°C, 60% relative humidity in a 12h/12h light-dark cycle on a standard cornmeal-based diet. Individual insertions are listed below. Detailed information on genotypes used in this study is listed in the Supplemental Genotypes and Statistics table.

| Insertion | Source | Reference |
| --- | --- | --- |
| R37H08-Gal4 | Kind gift of Y. Aso, HHMI, Janelia | FlyBase: FBti0191253 |
| P{UASp-Venus.GAP43}10 | Bloomington Drosophila Stock center | FlyBase: FBti0129930 |
| P{UAS-tdTom.S}2 | Bloomington Drosophila Stock center | FlyBase: FBti0145103 |
| P{ry[+t7.2]=lArB}rut[2080] | Kind gift of A. Fiala, Uni Goettingen | [49]<br>FlyBase: FBti0003267 |
| P{UAS-GCaMP3.homer} | Kind gift of A. Fiala, Uni Goettingen | [30] |
| MB247-Dα7::GFP | Lab stock collection | [29] |
| UAS-brp-short<sup>cherry</sup> | Lab stock collection | [29] |

### Behaviour

All experimental steps were performed at 23°C, 60% relative humidity using mixed populations of *Drosophila* males and females maintained in a 12h/12h light/dark cycle. Flies were collected 0-4d after eclosion, starved for 24 hours on wet paper tissue (Kimberly-Clark Worldwide Inc.) allowing for water uptake and then trained. In appetitive memory experiments ∼80 flies were first exposed to an odour (CS-) alone (2min in short- and 5 minutes in long-term memory experiments). After a 2min inter-stimulus pause flies were trained by receiving dry sucrose on filter paper (3M Chr, Whatman) paired with a second odour (CS+) (2min in short- and 5 minutes in long-term memory experiments). 5 minutes of sugar availability improved the survival of flies undergoing the LTM paradigm. In mock controls, all stimuli used in the associative conditioning experiment were presented separately. Flies were tested after 1min retention time for short-or after 24h retention time for long-term memory. During the 24h retention flies were deprived of food and maintained in tubes containing moist paper tissue. During the test flies were allowed to choose between CS+ and CS-odours in a T-maze for 2min. A preference index (PI) was then calculated as the ratio of the difference between the number of flies that chose the CS+ and those that chose the CS-odour and the total number of flies: 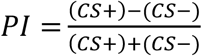. Odours used for conditioning were 11-cis vaccenyl acetate (Cayman Chemicals) 1:400 in 5% EtOH in PBS, geranyl acetate (Sigma Aldrich) 1:100 in 5% EtOH in PBS or 5% EtOH in PBS. EtOH was necessary to provide a food-related context to the starved flies (Figure S2B).

### De-novo protein synthesis inhibition

Immediately after training, flies were fed 35 mM cycloheximide (Sigma-Aldrich) [6] dissolved in 125mM sucrose and 0.01% carmine solution for 30min. The red dye carmine allowed confirming rapid drug uptake. A control group fed with 125mM sucrose and 0.01% carmine (Sigma-Aldrich) solution showed no learning defects.

### Immunohistochemistry

2-5 flies were randomly picked from conditioning experiments right before testing. Brains of females were dissected in cold phosphate-buffered saline (PBS) with 0.05% Triton and subsequently fixed in PBS containing 4% formaldehyde at RT for 50 min. After fixation brains were washed in PBS with 0.3% Triton before incubation overnight at 4°C with the following primary antibodies all diluted in PBS with 0.3% Triton: rabbit anti-RFP (1:2000; Rockland), rabbit anti-GFP (1:200; Life Technologies), mouse monoclonal anti-synapsin (3C11, 1:100; DSHB), mouse monoclonal anti-β-Galactosidase (1:200 Abcam). After washing, the brains were incubated with secondary antibodies in PBS containing 0.3% Triton for 4h at RT.

The secondary antibodies were Alexa Fluor568-conjugated goat anti-rabbit, Alexa Fluor488-conjugated goat anti-rabbit, Alexa Fluor568-conjugated goat anti-mouse, Alexa Fluor633-conjugated goat anti-mouse (all used 1:200 and from Life Technologies). Brains were mounted in Vectashield (Vector) and imaged with a laser scanning confocal microscope (LSM 780, Zeiss). For high resolution scans we used a C Plan-Apochromat 63x/1,4 Oil objective (Zeiss) with a voxel size of 0.09×0.09×0.25µm^3^ for quantitative analysis. Overviews of entire brains were taken with an LCI Plan-Apochromat 25x/0.8 objective (Zeiss) at a voxel size of 0.55×0.55×1µm^3^.

### Axon reconstruction

To analyse axon and bouton distribution in the calyx, membrane-tagged Venus was expressed in addition to the previously used markers under the control of a DA1-PN Gal4-driver line (*R37H08-GAL4, UAS-Gap43::Venus / MB247-Dα7::GFP, UAS-brp-short*^*cherry*^). Brains of 10 female flies per condition were immunolabelled with anti-synapsin antibodies (as above) and imaged. For PN axon reconstruction a high-resolution scan (0.09×0.09×0.25µm^3^, 63x NA1.4 oil immersion) of the right brain hemisphere of female flies was acquired with a confocal microscope. In addition, an overview scan used for registration was taken with a low magnification objective (25x; NA 0.8 multi-immersion). PN axon reconstruction was performed on the high-resolution scan of Venus signal in the trees toolbox available for Matlab [72]. In a second step tracings and high-resolution images were aligned to the registered calyx. For generation of a standard calyx with a volume of 37583 µm^3^ (Figure 3E, light green) the Dα7 signal of three registered calyces was averaged and reconstructed in Amira using the segmentation editor. Next, tracings were aligned to the registered overview scan in two steps. First, the iACT of the high-resolution image and of the registered brain in the Venus channel were aligned. Next, the calyx volume of the high-resolution calyx and the standard calyx went through a rigid registration performed in Amira. The alignment parameters were then applied to the axon reconstructions. Boutons were traced on the now registered high-resolution images with the landmark function in Amira. Bouton distribution inside the MBC was evaluated within a 3D grid of 10μm^3^ cubes.

### Two-photon *in vivo* calcium imaging

A mixed population of up to 4 d old *MB247-homer::GCaMP3* flies were starved for 18h at 22°C before appetitive conditioning with cVA (1:400, 5% EtOH in PBS), GA (1:100, 5% EtOH in PBS) was used as CS-. Starved untrained flies displayed no bias towards either of these odours at 24 hours. Flies used for imaging were randomly picked from the trained group right before testing. They were used for imaging only if the remaining flies from the same group had learned. For imaging, flies were briefly anesthetized on a peltier element at 4°C, placed into a custom-built imaging chamber (Figure 4A) and fixed using adhesive tape. The head capsule was opened under Ringer’s solution (5 mM HEPES, pH 7.4, 130 mM NaCl, 5 mM KCl, 2 mM CaCl_2_, 2 mM MgCl_2_). To minimize movement brains were stabilized with 1,5% low melting agarose (Thermo Scientific) in Ringer’s solution. Flies were imaged with a two-photon laser-scanning microscope (LaVision BioTec, TriM Scope II) equipped with an ultra-fast z-motor (PIFOC® Objective Scanner Systems 100µm range) and a Zeiss C-Apochromat 40x, 1.1 NA water –immersion objective. Two-photon images were analysed using Fiji/ImageJ [73]. GCaMP fluorescence was excited at 920 nm using a Ti:sapphire laser (Coherent Chameleon). A stack consisting of ∼ 10 optical sections was taken at 1Hz in approximately 0,26×0,26µm xy- and at 4µm z-resolution. Odours were applied with a constant humidified air stream (10ml/s) using a commercial device (Stimulus Controller CS 55, Ockenfels SYNTECH GbmH) triggered 5s after acquisition of the 1st frame by a multifunction I/O module (NI USB-6008), which was controlled by Matlab (Data Acquisition Toolbox). To record DA1 neurons specific responses, *UAS-tdTomato; R37H08-GAL4, MB247-Homer::GCaMP3* flies were anesthetized on ice, positioned in a polycarbonate imaging chamber ([74]; gift from S. Tomchik), and immobilized using Myristic Acid (Sigma-Aldrich). To allow optical access to the Calyx, a small window was opened through the head capsule under Ringer’s solution. Two-photon microscopy was conducted as described above.

### Two-photon image data processing

The time series was processed with a custom Fiji/ ImageJ macro and corrected for small x/y shifts with the StackReg plug-in [75]. A grid (ROIs, side length 5 μm) was assigned for each optical slide of a stack covering the entire calyx. Intensity tables of each square of the grid were exported to Microsoft Excel and the ΔF/F was calculated. The baseline (F_0_) was set by averaging the intensities within each ROI of the 5 frames prior to odour stimulation. ROIs were regarded as responsive, if their normalized ΔF/F% throughout the first 2s of odour application exceeded 3x the standard deviation of the F_0_ of the same ROI in the 5s (=5 images) before odour stimulation. ROIs below that threshold were assigned into the category “unresponsive”. ROIs calcium responses higher than the threshold were further subdivided into three categories. The first category was “Carrier” (responsive to both, 5% EtOH and 1:400 cVA, 5% EtOH). The second category was “cVA” (responsive only to 1:400 cVA, 5% EtOH and not to 5% EtOH) or “EtOH” (response only to 5% EtOH application and not to 1:400 cVA, 5% EtOH). To analyse DA1 boutons responses to 5% EtOH and 1:400 cVA+ 5% EtOH were recorded from naïve *UAS-tdTomato; R37H08-GAL4, MB247-Homer::GCaMP3* flies, exported to Fiji/ ImageJ and ROIs were manually drawn around DA1 boutons based on the tdTomato fluorescence. Intensity values of each ROI were transferred to Microsoft Excel and ΔF/F values were calculated using the average of the first 5 frames prior to odour stimulation as baseline (F_0_). Responsive ROIs were defined as above. These results were age, gender and sequence independent as presenting the odours in a different order did not change the results of the analysis. Calcium traces were generated in Prism 7 (GraphPad Software).

### Statistics

Statistical analyses were performed with Prism7.01 software (GraphPad). All data were tested for normality (D’Agostino & Pearson omnibus normality test) and homogeneity of variances (F-test). Comparisons of normally distributed data were tested by a one-sample t test, a two-sample t test or one-way analysis of variance (ANOVA) followed by planned, pairwise multiple-comparison tests with adjusted p-values (Bonferroni), as shown in the Genotypes and Statistics table. Definition of statistical significance was set to <0.05. Asterisks denote * *p*<0.05; ** *p*<0.01; \*\*\**p*<0.001; **** *p*<0.0001; n.s. not significant.

