## Supplementary Figures for "Circuit reorganization in the *Drosophila* mushroom body calyx accompanies memory consolidation"

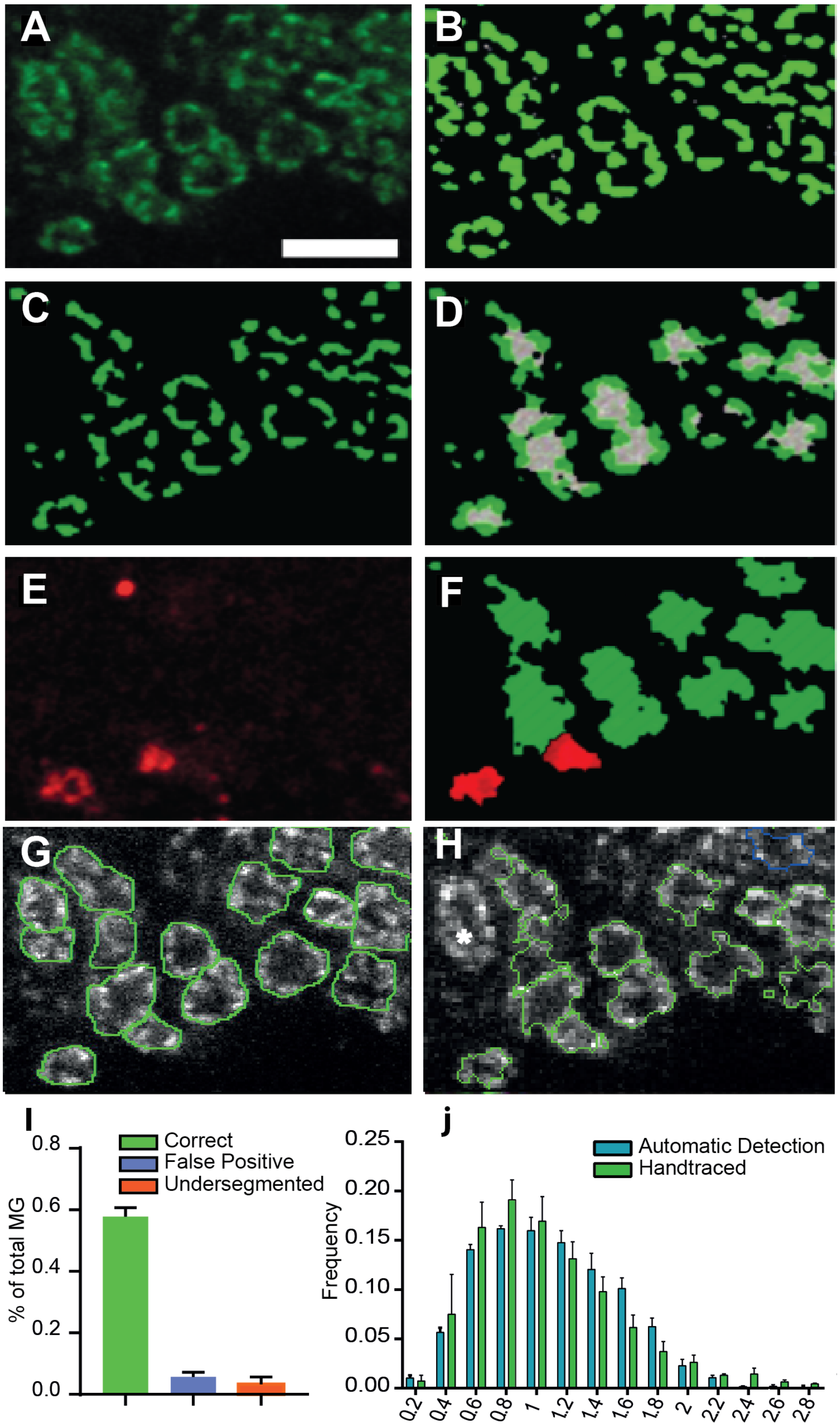

**Figure S1| Automated identification and reconstruction of microglomeruli (Related to Figure 1)**

**(A)** Example of a small area from a typical optical section used for the automated detection. Only the green channel containing the Dα7 signal was used for identification of MGs. An anisotropic filter was applied to the original image and contrast was enhanced. Scale bar = 10μm. **(B)** Initial segmentation of the entire image was performed by grouping pixels with similar grey values into individual objects (green). **(C)** A membership function assigned by applying a histogram shape-based threshold on the brightest objects of this contrast map as candidate objects for MG rings (dark green). **(D)** A second threshold was set to assign seed points for MG lumen within darkest areas surrounded by MG ring candidates. These seed lumen candidate objects grew in a watershed analysis 1 pixel for 5 cycles in 3D into dark areas or until a MG ring candidate object was reached. Lumen candidate objects were classified as “real lumen” (grey) by a fuzzy classification approach depending on lumen candidate volume, their elliptic fit (or roundness) and their relative border with ring candidate objects. Final MG rings (light green) were finally detected in 3D by watersheding into bright areas using the lumens as seed points. **(E)** The red channel containing the Brp-short$^{cherry}$ signal was applied to create objects representing labelled active zones. **(F)** MGs identified in the green channel image that colocalized with the independently generated objects in the red channel image were classified as R37H08-positive (DA1- MGs; red). MGs that did not colocalize with Brp-short$^{cherry}$ signal were displayed in green. **(G)** To evaluate the performance of the routine, all MGs within 4 MBCs were traced manually (green rings) and were overlaid in a custom Matlab script with automatic reconstruction. **(H)** Result of the comparison of the automated and the manual reconstruction shown in (G). MGs detected in the manual reconstruction and in the automated reconstruction are displayed in green (*“correct”*). MGs only detected in the automated reconstruction (“*false-positive”*) are displayed in blue. Additional possible errors of the reconstruction were “*false-negative”,* if a MG was detected only in the manual reconstruction, or “*undersegmented*”, if multiple MG were fused in the automated reconstruction compared to the manual reconstructions. **(I)** Statistical evaluation of the performance of the automated reconstruction. Compared to the manual reconstruction the automated method detected 59% of all MG. ~3.7% were false positive and ~2.4% undersegmented. **(J)** Comparison of the size distribution of MG detected manually (green) or by the software (blue) in the same optical sections (n of calyces = 4). Comparison of the relative size distribution frequencies with Wilcoxon Matched Pairs Test revealed no significant difference between manual and automated MG identification (mean $p>0.5$).

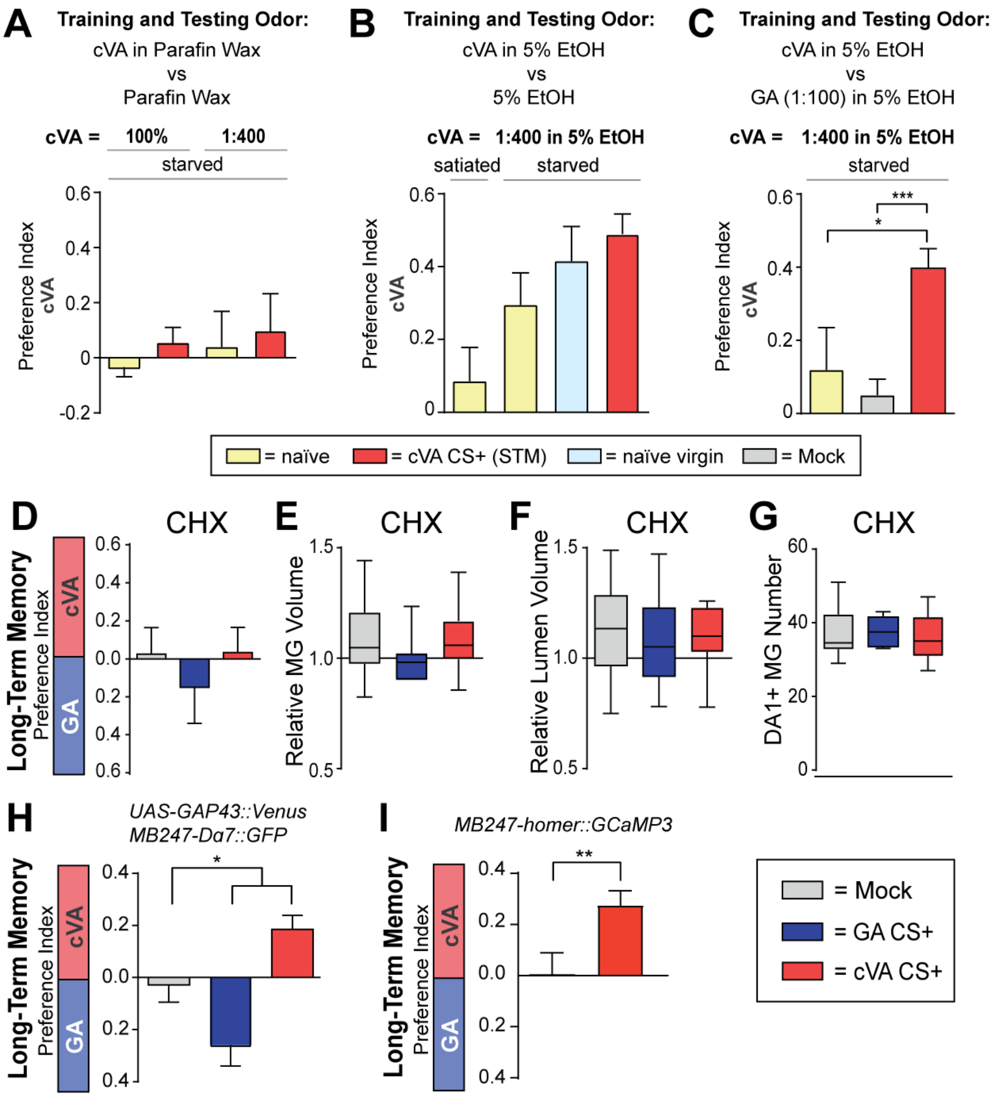


**Figure S2| Establishment of appetitive conditioning with cVA, suppression of anatomical modifications after pharmacological blocking of long-term memory (Related to Figure 2) and learning scores of individual genotypes (Related to Figures 3, 4)**

**(A)** Appetitive STM conditioning using pure cVA or cVA diluted 1:400 in paraffin wax induced preference scores towards cVA that were similar to those of naïve flies. **(B)** The pheromone cVA was attractive for naïve flies when applied in a food context (choice: 1:400 cVA + 5% EtOH versus 5% EtOH). Attraction was stronger if flies were starved, if virgin females were tested or after appetitive conditioning using cVA (1:400 in 5% EtOH) as CS+ and the carrier (5% EtOH) as CS-. **(C)** Comparison of preference scores towards cVA of flies that were confronted with the choice between cVA (1:400 in 5% EtOH) and GA (1:100 in 5% EtOH). Flies that had been trained in an appetitive paradigm with cVA (1:400 in 5% EtOH) as CS+ and GA (1:100 in 5% EtOH) as CS- displayed higher preference scores towards cVA in comparison to mock-trained flies or starved naïve flies, indicating that flies can learn to associate cVA with a reward in these conditions **(D)** Pharmacological suppression of LTM by feeding *R37H08-GAL4/ UAS-brp-short*$^{cherry}$*, MB247-Dα7::GFP* flies 50 mM cycloheximide (CHX) in 125mM sucrose solution for 30 min after training. Subsequently flies were re-starved for 24h before testing (p > 0.05, n = 8–11). **(E-G)** Suppression of the structural modifications in DA1- MGs in cVA CS+ flies after LTM block by CHX application. ($p > 0.05$). n = 7–17. Compare to Figure 2G-I. **(H)** Appetitive LTM scores of *R37H08-Gal4, UAS-GAP43::Venus /MB247-Dα7::GFP, UAS-brp-short*$^{cherry}$ flies (*p < 0.5; n= 23- 32), used in Figure 3A-H. **(I)** Appetitive LTM scores of *MB247-homer::GCaMP3* flies ($p < 0.01$; t test; n = 32-34), used in Figure 4A-F and in Figure S3.

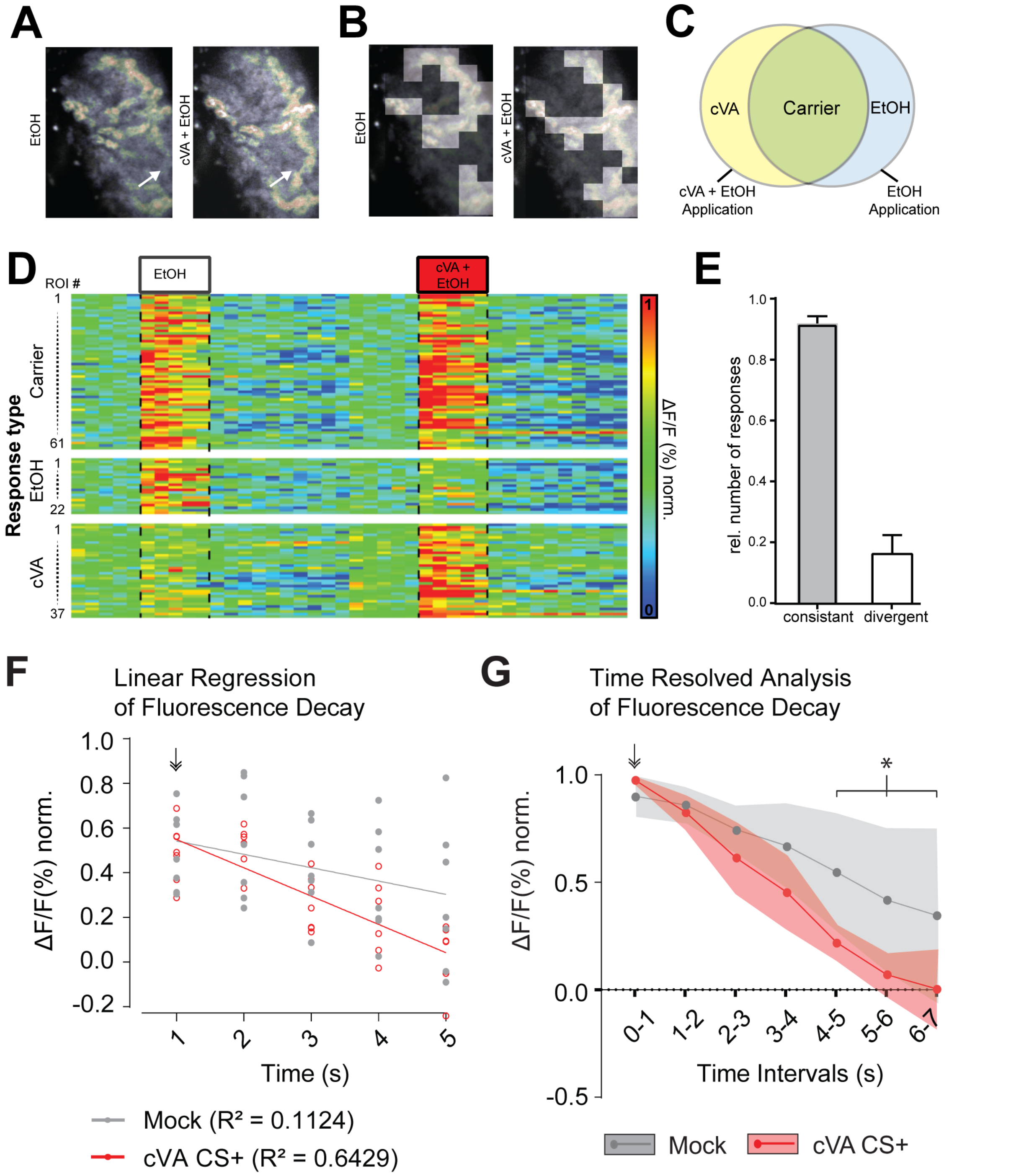


**Figure S3| Classification of calcium responses in the MBC (Related to Figure 4)**

**(A)** Representative optical section of a time z-stack series consisting of 30 cycles with 10 sections per stack at a frame rate of 1 Hz. The odour was applied for 5s after 5 cycles. With a typical calycal diameter along the dorsal/ventral axis of 35µm and an average MG diameter of 5µm this imaging settings reliably captured calcium dynamics of the entire MBC. The grey scale image was created by averaging images from one optical plane over the last 15 acquisition cycles after odour application. False color-coded images were created by subtracting the background (generated by averaging 5 images preceding the odour application) from the first two averaged images during odour application. White arrows point to areas only responsive to cVA. **(B)** Single optical planes were overlaid with a grid of 5x5 $\mu m^2$ meshes (ROIs). White squares represent ROIs, which were classified as responsive to odour stimulation. For classification the mean response as $\Delta F/F\%$ during the first 2s of odour application was calculated for each ROI. ROIs were classified as odour responsive, if the mean $\Delta F/F\%$ during the first 2s of odour application was greater than 3x the standard deviation of the $\Delta F/F\%$ during the 5s preceding odour stimulation. **(C)** Schematic of ROI classification strategy. ROIs were classified as responsive to "cVA" if they displayed an above-threshold response to cVA (1:400, 5% EtOH), but not to EtOH only (5% EtOH). "Carrier" ROIs responded to both stimuli and "EtOH" only to EtOH (5% EtOH), but not to cVA (1:400, 5% EtOH). **(D)** Temporal dynamics of fluorescence changes to odour stimulation within a single MBC. Each lane of the matrix represents a single ROI identified as responsive towards the carrier (top matrix), EtOH only (mid matrix) or cVA (bottom matrix) as in (C). Each column of the matrix represents 1s. White and red boxes and dashed lines represent 5s of odour stimulation with 5% EtOH or cVA (1:400, 5% EtOH), respectively. Out of 120 responsive ROIs in one MBC, 61 ROIs were classified as responsive to the carrier, 22 to EtOH and 37 were specific to cVA. **(E)** Number of consistent responses in the MBC after repetitive odour stimulation with 5% EtOH. After two consecutive odour stimulations ~90% of the responsive areas kept consistent compared to a previous stimulation ($n = 6$). The fraction of areas only responsive during 2nd odour stimulation and classified as divergent was ~16%. **(F)** Comparison of calcium signal decay in KC dendrites by linear regression analysis. From its peak after the start (arrow) of five seconds stimulation with cVA, calcium response decays more rapidly in cVA CS+ trained flies than in mock trained flies. A linear fit described the decrease of GCaMP fluorescence well in cVA CS+ trained flies, whereas decay was less homogenous in flies from the mock trained group. ($p < 0.05$; $n = 7$). **(G)** Time resolved analysis of fluorescence decay of GCaMP after the initial peak. During the initial two seconds of cVA stimulation (arrow) average responses in KC dendrites reached their peak in each condition. The calcium response was undistinguishable between the two groups until 3-4 s of stimulation, but it decayed more rapidly from 4-5 s on in the cVA CS+ flies. ($p < 0.05$; $n = 7$)

| Connection to DA1-PN bouton | Cell type | Number of cells | Number of synaptic connections |
|---|---|---|---|
| Postsynaptic to the DA1-PN bouton | KC αβp | 0 | 0 |
| | KC αβc | 1 | 12 |
| | KC αβs | 3 | 11-21 |
| | KC α’β’ap | 1 | 25 |
| | KC α’β’m | 1 | 8 |
| | KC γmain | 8 | 12-24 |
| | KC γd | 0 | 0 |
| | APL | 1 | 2 |
| | MB-C1 | 2 | 1-3 |
| Presynaptic to the DA1-PN bouton | KC γmain | 2 | 1 |
| | APL | 1 | 3 |
| | MB-C1 | 1 | 1 |

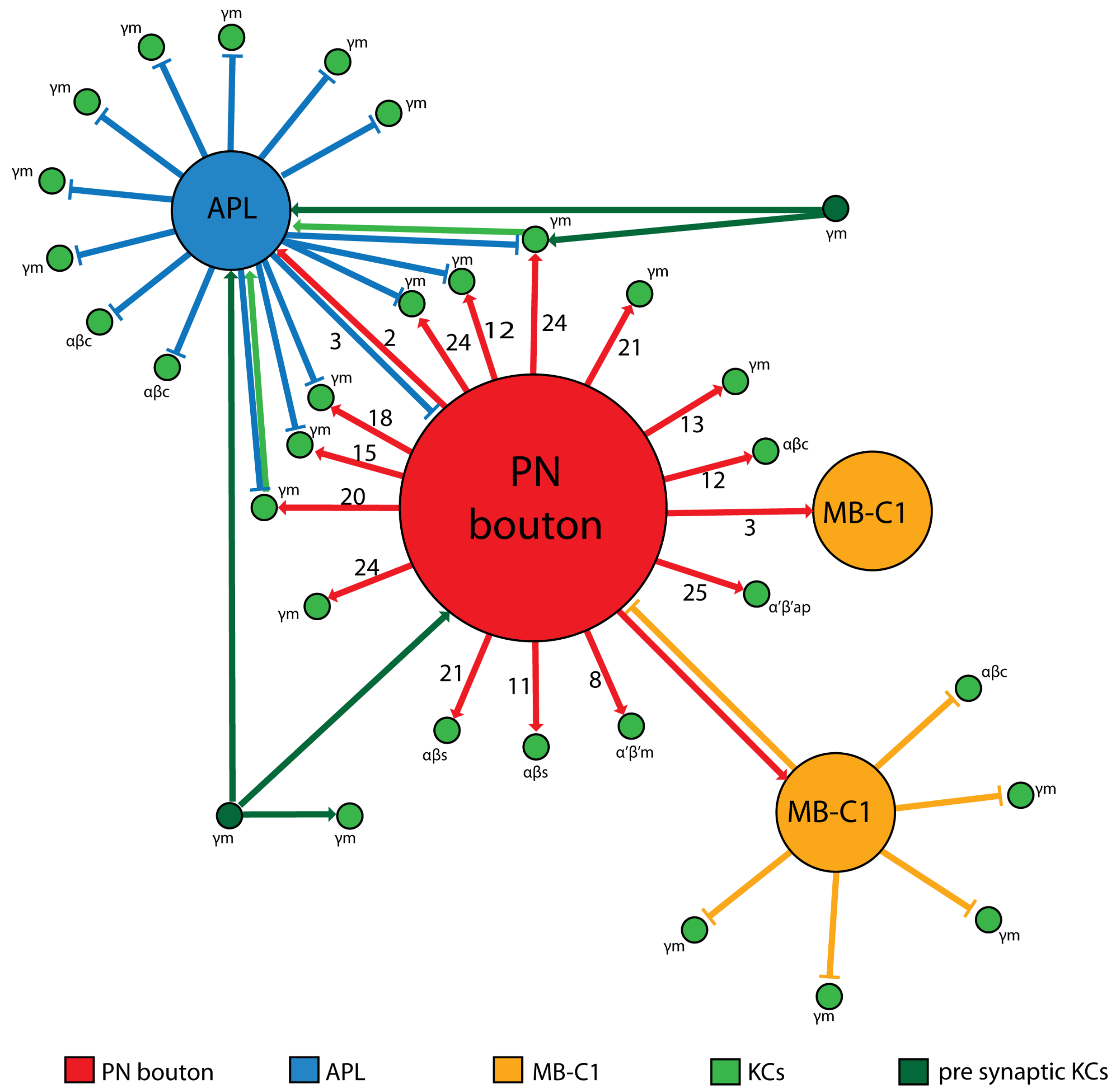

**Supplementary Table1| Neurons contributing to a DA1-PN microglomerulus and their synaptic connections (Related to Figure 1)**

Top: Table representing the identified direct synaptic contacts between the DA1-PN bouton (see Figure 1) and other neurons, revealed from reconstructions in the FAFB EM dataset [1]. Bottom: Scheme of the MG network of the DA1-PN bouton (see Figure 1) reconstructed in the FAFB EM dataset. If synaptic connections are more than one, their number is indicated along the arrows. The network consists of a DA1-PN bouton (red), surrounded by the APL neuron (blue), two MB-C1 neurons (orange), 14 postsynaptic KC claws and two additional KCs (dark green), which are presynaptic to the PN bouton and to APL. APL and MB-C1 form polyadic synapses with the PN bouton including KCs. Some of these are not postsynaptic to the PN bouton and are therefore placed in the scheme around the respective APL or MB-C1 neuron.

**Movie 1| 3D reconstruction of a DA1-PN microglomerulus (Related to Figure 1)**

Reconstruction of a microglomerulus from EM serial sections derived from FAFB data set [1]. The movie shows a DA1-PN bouton (red) and all profiles that are directly pre- or postsynaptic to it. The DA1-PN bouton is surrounded by the claws of 14 KCs of five subtypes (different green shades according to Figure 1D, E). Three additional neurons contribute to this microgolomerulus: APL (blue), which is pre- and postsynaptic to the PN bouton, and both MB-C1 neurons (yellow). Finally, two γmain KCs (dark green) form presynaptic connections with the bouton. See also Supplementary Table 1

### Genotypes and statistics

| Figure | Group code | n | Fly strain | tested groups | test type | (adjusted) p value |
|---|---|---|---|---|---|---|
| Fig 1C | | 7 | *w-; UAS-tdTomato; R37H08-Gal4/MB247-homer::GCaMP3* | | | |
| Fig 1F-G | | | *w-; ;R37H08-Gal4/MB247-Dα7::GFP,UAS-brpshort::Cherry* | | | |
| Fig 2B | 1 | 19 | *w-; ;R37H08-Gal4/MB247-Dα7::GFP,UAS-brpshort::Cherry* | all | one-way ANOVA | <0,0001 |
| | 2 | 24 | | 1 vs 2 | mean of group 2 and 3 compared with control column 1 | <0,0001 |
| | 3 | 25 | | 1 vs 3 | ” | <0,0001 |
| Fig 2C | | | *w-; ;R37H08-Gal4/MB247-Dα7::GFP,UAS-brpshort::Cherry* | all | one-way ANOVA | 0,8999 |
| | 1 | 14 | | 1 vs 2 | post-hoc test with Bonferroni correction | 0.2816 |
| | 2 | 15 | | 1 vs 3 | " | 0.4576 |
| | 3 | 20 | | 2 vs 3 | " | 0.1708 |
| Fig 2D | | | | all | one-way ANOVA | P=0,9105 |
| | 1 | 14 | *w-; ;R37H08-Gal4/MB247-Dα7::GFP,UAS-brpshort::Cherry* | 1 vs 2 | post-hoc test with Bonferroni correction | 0.1548 |
| | 2 | 15 | | 1 vs 3 | " | 0.2575 |
| | 3 | 18 | | 2 vs 3 | " | 0.427 |
| Fig 2E | | | *w-; ;R37H08-Gal4/MB247-Dα7::GFP,UAS-brpshort::Cherry* | all | one-way ANOVA | 0,7791 |
| | 1 | 11 | | 1 vs 2 | post-hoc test with Bonferroni correction | 0.5479 |
| | 2 | 11 | | 1 vs 3 | " | 0.07376 |
| | 3 | 17 | | 2 vs 3 | " | 0.6775 |
| Fig 2F | 1 | 17 | *w-; ;R37H08-Gal4/MB247-Dα7::GFP,UAS-brpshort::Cherry* | all | one-way ANOVA | 0.9406 |
| | 2 | 14 | | 1 vs 2 | post-hoc test with Bonferroni correction | 0.0148 |
| | 3 | 19 | | 1 vs 3 | " | 0.0183 |
| Fig 2G | | | *w-; ;R37H08-Gal4/MB247-Dα7::GFP,UAS-brpshort::Cherry* | all | one-way ANOVA | 0,0309 |
| | 1 | 17 | | 1 vs 2 | post-hoc test with Bonferroni correction | 0.8507 |
| | 2 | 14 | | 1 vs 3 | " | 0.0137 |
| | 2 | 19 | | 2 vs 3 | " | 0.0366 |
| Fig 2H | | | *w-; ;R37H08-Gal4/MB247-Dα7::GFP,UAS-brpshort::Cherry* | all | one-way ANOVA | 0,0030 |
| | 1 | 17 | | 1 vs 2 | post-hoc test with Bonferroni correction | 0.3292 |

| | | | | | | |
|---|---|---|---|---|---|---|
| | 2 | 14 | | 1 vs 3 | " | 0.0421 |
| | 2 | 19 | | 2 vs 3 | " | 0.0024 |
| Fig 2I | | | *w-; ;R37H08-Gal4/MB247-Dα7::GFP,UAS-brpshort::Cherry* | all | one-way ANOVA | 0.0066 |
| | 1 | 24 | | 1 vs 2 | post-hoc test with Bonferroni correction | >0,9999 |
| | 2 | 18 | | 1 vs 3 | " | 0.012 |
| | 3 | 21 | | 2 vs 3 | " | 0.0287 |
| Fig 2J | 1 | 14 | *rut2080; ;R37H08-Gal4/MB247-Dα7::GFP,UAS-brpshort::Cherry* | all | one-way ANOVA | 0.5896 |
| | 2 | 18 | " | 1 vs 2 | mean of group 2 and 3 compared with control column 1 | >0,9999 |
| | 3 | 17 | " | 1 vs 3 | " | >0,9999 |
| Fig 2K | 1 | 21 | *rut2080; ;R37H08-Gal4/MB247-Dα7::GFP,UAS-brpshort::Cherry* | all | one-way ANOVA | 0.4755 |
| | 2 | 17 | " | 1 vs 2 | post-hoc test with Bonferroni correction | 0.7165 |
| | 3 | 20 | " | 1 vs 3 | " | >0,9999 |
| | | | | 2 vs 3 | " | >0,9999 |
| Fig 2L | 1 | 21 | *rut2080; ;R37H08-Gal4/MB247-Dα7::GFP,UAS-brpshort::Cherry* | all | one-way ANOVA | 0.7071 |
| | 2 | 17 | " | 1 vs 2 | post-hoc test with Bonferroni correction | >0,9999 |
| | 3 | 20 | " | 1 vs 3 | " | >0,9999 |
| | | | | 2 vs 3 | " | >0,9999 |
| Fig 2M | 1 | 18 | *rut2080; ;R37H08-Gal4/MB247-Dα7::GFP,UAS-brpshort::Cherry* | all | one-way ANOVA | 0.8146 |
| | 2 | 16 | " | 1 vs 2 | post-hoc test with Bonferroni correction | >0,9999 |
| | 3 | 13 | " | 1 vs 3 | " | >0,9999 |
| | | | | 2 vs 3 | " | >0,9999 |
| Fig 3A-E | | | *w-; ;R37H08-Gal4, UAS-Gap43::Venus/MB247-Dα7::GFP,UAS-brpshort::Cherry* | | | |
| Fig 3F | 1 | 10 | *w-; ;R37H08-Gal4, UAS-Gap43::Venus/MB247-Dα7::GFP,UAS-brpshort::Cherry* | | one-way ANOVA | <0,0001 |
| | 2 | 10 | | 1 vs 2 | post-hoc test with Bonferroni correction | <0,0001 |
| | 3 | 10 | | 1 vs 3 | " | <0,0001 |
| | | | | 1 vs 4 | " | <0,0001 |
| | | | | 2 vs 3 | " | 0.7129 |
| | | | | 2 vs 4 | " | >0,9999 |

| | | | | | | |
|---|---|---|---|---|---|---|
| | | | | 3 vs 4 | " | 0.4852 |
| Fig 3G | 1 | 10 | *w-; ;R37H08-Gal4, UAS-Gap43::Venus/MB247-Dα7::GFP,UAS-brpshort::Cherry* | all | one-way ANOVA | 0.0025 |
| | 2 | 10 | | 1 vs 2 | post-hoc test with Bonferroni correction | 0.1104 |
| | 3 | 10 | | 1 vs 3 | " | 0.0018 |
| | | | | 2 vs 3 | " | 0.4569 |
| Fig 3H | 1 | 10 | *w-; ;R37H08-Gal4, UAS-Gap43::Venus/MB247-Dα7::GFP,UAS-brpshort::Cherry* | all | one-way ANOVA | 0.0217 |
| | 2 | 10 | | 1 vs 2 | post-hoc test with Bonferroni correction | 0.9985 |
| | 3 | 10 | | 1 vs 3 | " | 0.0471 |
| | | | | 2 vs 3 | " | 0.0382 |
| Fig 4B-C | | | *w-; ;mb247-homer-GCamp3/TM3* | | | |
| Fig 4D | 1 | 7 | *w-; ;mb247-homer-GCamp3/TM3* | 1 vs 2 | t test | 0.0191 |
| | 2 | 7 | " | | " | |
| Fig 4E-F | | | *w-; ;mb247-homer-GCamp3/TM3* | | | |
| Fig S1A-H | | | *w-; ;R37H08-Gal4/MB247-Dα7::GFP,UAS-brpshort::Cherry* | | | |
| Fig S1I-J | 1 | 4 | *w-; ;R37H08-Gal4/MB247-Dα7::GFP,UAS-brpshort::Cherry (automated)* | 1 vs 2 | Kolmogorov-Smirnov test | 0.0382 |
| | 2 | 4 | *w-; ;R37H08-Gal4/MB247-Dα7::GFP,UAS-brpshort::Cherry (hand traced)* | | " | 0.5085 |
| Fig S2A | 1 | 12 | *Canton S* | 1 vs 2 | t test | 0.2344 |
| | 2 | 16 | " | 3 vs 4 | " | 0.7574 |
| | 3 | 8 | " | | | |
| | 4 | 8 | " | | | |
| Fig S2B | 1 | 9 | *Canton S* | 2 vs 3 vs 4 | one-way ANOVA | 0.1634 |
| | 2 | 15 | " | 2 vs 3 | post-hoc test with Bonferroni correction | 0.9898 |
| | 3 | 9 | " | 2 vs 4 | " | 0.1767 |
| | 4 | 19 | " | 3 vs 4 | " | >0,9999 |
| Fig S2C | 1 | | *Canton S* | all | one-way ANOVA | |
| | 2 | | " | 2 vs 3 | post-hoc test with Bonferroni correction | >0,9999 |
| | 3 | | " | 2 vs 4 | " | 0.0214 |
| | | | | 3 vs 4 | " | 0.0007 |

| | | | | | | |
|---|---|---|---|---|---|---|
| Fig S2D | 1 | 8 | *w-; ;R37H08-Gal4/MB247-Dα7::GFP,UAS-brpshort::Cherry* | all | one-way ANOVA | 0.6831 |
| | 2 | 4 | " | 1 vs 2 | mean of group 2 and 3 compared with control column 1 | >0,9999 |
| | 3 | 11 | " | 1 vs 3 | " | >0,9999 |
| Fig S2E | 1 | 10 | *w-; ;R37H08-Gal4/MB247-Dα7::GFP,UAS-brpshort::Cherry* | all | one-way ANOVA | 0.4207 |
| | 2 | 8 | " | 1 vs 2 | post-hoc test with Bonferroni correction | 0.6737 |
| | 3 | 17 | " | 1 vs 3 | " | >0,9999 |
| | | | | 2 vs 3 | " | 0.7306 |
| Fig S2F | 1 | 10 | *w-; ;R37H08-Gal4/MB247-Dα7::GFP,UAS-brpshort::Cherry* | all | one-way ANOVA | 0.7071 |
| | 2 | 8 | " | 1 vs 2 | post-hoc test with Bonferroni correction | >0,9999 |
| | 3 | 17 | " | 1 vs 3 | " | >0,9999 |
| | | | | 2 vs 3 | " | >0,9999 |
| Fig S2G | 1 | 10 | *w-; ;R37H08-Gal4/MB247-Dα7::GFP,UAS-brpshort::Cherry* | all | one-way ANOVA | 0.8263 |
| | 2 | 8 | " | 1 vs 2 | post-hoc test with Bonferroni correction | >0,9999 |
| | 3 | 17 | " | 1 vs 3 | " | >0,9999 |
| | | | | 2 vs 3 | " | >0,9999 |
| Fig S2H | 1 | 24 | *w-; ;R37H08-Gal4, UAS-Gap43::Venus/MB247-Dα7::GFP,UAS-brpshort::Cherry* | all | one-way ANOVA | <0,0001 |
| | 2 | 23 | | 1 vs 2 | post-hoc test with Bonferroni correction | 0.0413 |
| | 3 | 32 | | 1 vs 3 | " | 0.0413 |
| Fig S2I | 1 | 32 | *w-; ;mb247-homer-GCamp3/TM3* | 1 vs 2 | t test | 0.0097 |
| | 2 | 34 | " | | " | |
| Fig S3A-E | | | *w-; ;mb247-homer-GCamp3/TM3* | | | |
| Fig S3F | 1 | 7 | *w-; ;mb247-homer-GCamp3/TM3 (cVA CS+; red)* | 1 vs 2 | Linear Regression analysis of slopes | 0.0433 |
| | 2 | 7 | *w-; ;mb247-homer-GCamp3/TM3 (mock; grey)* | | | |
| Fig S3G | 1 | 7 | *w-; ;mb247-homer-GCamp3/TM3 (cVA CS+;red)* | 1st interval | multiple t test with | 0.9964595 |
| | 2 | 7 | " | 2nd interval | | 0.9998686 |
| | | | | 3rd interval | | 0.8253874 |
| | | | | 4th interval | | 0.3098746 |
| | | | | 5th interval | | 0.0272125 |

| | | |
|---|---|---|
| | 6th interval | 0.0162665 |
| | 7th interval | 0.0187378 |

1. Zheng, Z., Lauritzen, J.S., Perlman, E., Robinson, C.G., Nichols, M., Milkie, D., Torrens, O., Price, J., Fisher, C.B., Sharifi, N., et al. (2018). A Complete Electron Microscopy Volume of the Brain of Adult Drosophila melanogaster. Cell *174*, 730-743 e722.
